# RibonucleaseH1-targeted R-loops in surface antigen gene expression sites can direct trypanosome immune evasion

**DOI:** 10.1101/361451

**Authors:** Emma Briggs, Kathryn Crouch, Leandro Lemgruber, Craig Lapsley, Richard McCulloch

**Affiliations:** The Wellcome Centre for Molecular Parasitology, University of Glasgow, College of Medical, Veterinary and Life Sciences, Institute of Infection, Immunity and Inflammation, Sir Graeme Davies Building, 120 University Place, Glasgow, G12 8TA, U.K

**Keywords:** antigenic variation, trypanosome, Variant Surface Glycoprotein, R-loop, RNase H1, DNA replication, transcription

## Abstract

Switching of the Variant Surface Glycoprotein (VSG) in *Trypanosoma brucei* provides a crucial host immune evasion strategy that is catalysed both by transcription and recombination reactions, each operating within specialised telomeric VSG expression sites (ES). VSG switching is likely triggered by events focused on the single actively transcribed ES, from a repertoire of around 15, but the nature of such events is unclear. Here we show that RNA-DNA hybrids, called R-loops, form preferentially within sequences termed the 70 bp repeats in the actively transcribed ES, but spread throughout the active and inactive ES in the absence of RNase H1, which degrades R-loops. Loss of RNase H1 also leads to increased levels of VSG coat switching and replication-associated genome damage, some of which accumulates within the active ES. This work indicates VSG ES architecture elicits R-loop formation, and that these RNA-DNA hybrids connect *T. brucei* immune evasion by transcription and recombination.

**Author summary:** All pathogens must survive eradication by the host immune response in order to continue infections and be passed on to a new host. Changes in the proteins expressed on the surface of the pathogen, or on the surface of the cells the pathogen infects, is a widely used strategy to escape immune elimination. Understanding how this survival strategy, termed antigenic variation, operates in any pathogen is critical, both to understand interaction between the pathogen and host and disease progression. A key event in antigenic variation is the initiation of the change in expression of the surface protein gene, though how this occurs has been detailed in very few pathogens. Here we examine how changes in expression of the surface coat of the African trypanosome, which causes sleeping sickness disease, are initiated. We reveal that specialised nucleic acid structures, termed R-loops, form around the expressed trypanosome surface protein gene and increase in abundance after mutation of an enzyme that removes them, leading to increased changes in the surface coat in trypanosome cells that are dividing. We therefore shed light on the earliest acting events in trypanosome antigenic variation.

## Introduction

The genome provides the blueprint for life and is normally protected from rapid content change by high fidelity DNA replication and a range of repair pathways. However, strategies for elevated rates of genome variation have evolved, some of which are genome-wide, such as in developmental chromosome fragmentation in ciliates (1) and chromosome and gene copy number variation during *Leishmania* growth (2, 3). More commonly, enhanced genome change is more localised and caused by deliberate lesion generation, such as during yeast mating type switching, which is induced by HO endonuclease-mediated cleavage in *Saccharomyces cerevisiae* (4) and locus-directed replication stalling in *Schizosaccharomyces pombe* (5). Rearrangements to generate mature receptors and antibodies expressed by T and B cells (6) occur throughout mammalian immune genes and are generated by RAG endonuclease-catalysed DNA breaks (6) or transcription-linked base modification (7). Reflecting the diversity in routes capable of initiating genome change, homologous recombination (HR), non-homologous end-joining and microhomology-mediated end-joining repair reactions have been implicated in the catalysis of these different reactions. Antigenic variation is a very widespread pathogen survival strategy, involving stochastic switches in surface antigens to thwart host adaptive immunity (8), and locus-directed gene rearrangement is a common route for the differing reactions used in bacteria, fungi and protists (9, 10). Since antigenic variation impedes vaccination, it is important to understand the potentially pathogen-specific events that initiate surface antigen gene switching. However, only in the bacteria *Neisseria gonorrhoeae* is initiation well understood; here, HR catalyses movement of silent, non-functional *pilS* genes into a *pilE* expression locus via transcription-induced guanine quadruplex formation (11). In no other pathogen has the initiation event(s) during antigenic variation been resolved in this detail.

Antigenic variation in *T. brucei* displays remarkable mechanistic complexity, since it involves both HR-directed rearrangement and transcriptional control of *VSG* genes. In any cell a single *VSG* is expressed by RNA Polymerase (Pol) I transcription of one of the multiple telomeric ES (12, 13). In each ES the *VSG* is proximal to the telomere and is co-transcribed with multiple *ESAGs* (expression site associated genes) (14). Invariably, the *VSG* and *ESAGs* are separated by stretches of 70 bp repeats, sequences also found upstream of thousands of further *VSGs* in the *T. brucei* genome (15-17). To execute a VSG coat switch, *T. brucei* can use HR to move one of around 1000 silent subtelomeric *VSGs* into the active VSG ES, displacing the resident VSG, or silence transcription from the active VSG ES and activate transcription from one of the silent VSG ES. Though a wide range of factors have been described that influence singular ES expression (12, 18-21) and execute VSG recombination (22), initiation of VSG switching remains poorly understood. Targeting yeast I-SceI endonuclease activity to the active VSG ES elicits *VSG* recombination (23, 24), but no endogenous ES-focused endonuclease has been described. Impaired telomere protection results in VSG ES breaks (25, 26) and critically short telomeres have been associated with increased VSG switching (27), but how such processes act in unperturbed cells is unknown. Finally, DNA replication mapping suggests that the active VSG ES, uniquely amongst these telomeric loci, is subject to early replication in mammal-infective (bloodstream form, BSF) parasite cells (28), but how this extrapolates to VSG switch initiation is unclear (29).

R-loops are stable RNA-DNA hybrids that form within the DNA helix, displacing single-stranded DNA. Though R-loops can arise from transcription, active roles are emerging in an range of genomic processes (30), including replication initiation (31, 32) and arrest (33), transcription activation and termination (30), telomere homeostasis (26, 34) and chromatin formation (35, 36). In addition, R-loops can lead to genome instability and mutation (37-39). Two distinct ribonuclease H enzymes, RNase H1 and RNase H2, are found in eukaryotes and can degrade RNA within R-loops (40). Here, we explore R-loop distribution in the VSG ES of both wildtype BSF *T. brucei* and in null mutants that lack a homologue of RNase H1. We show that R-loops accumulate throughout all VSG ES in the absence of the RNase H enzyme, indicating RNA-DNA hybrids form in these transcription sites and are normally resolved by removing the RNA. Loss of the RNase H results in elevated levels of replication-associated damage and leads to increased VSG switching, suggesting a model for the events that initiate antigenic variation in *T. brucei.*

## Results

### *T. brucei* encodes a non-essential, nuclear RNaseH1

*T. brucei* encodes an RNase H1 homologue (TbRH1) that has been predicted to be nuclear by fusing an N-terminal fragment to GFP (41). By expressing full-length TbRH1 C-terminally fused to 12 copies of the myc epitope (Fig.S1A), from its own locus, we confirmed nuclear localisation (Fig.1A). Expression of TbRH1-myc was constitutive throughout all discernible cell cycle stages of BSF *T. brucei* cells and did not display any obvious sub-nuclear localisation (Fig.1B), though increased signal appeared present in cells undergoing nuclear replication (Fig.S1B). By integration of *TbRH1* targeting constructs (Fig.S2A) we generated heterozygous (+/-) and then homozygous (-/-) TbRH1 mutants (Fig.S2B). No growth perturbation (Fig.1C) or alteration in cell cycle stage distribution (Fig.1D) was apparent in the mutants, indicating TbRH1 does not provide essential genome functions (at least in culture), unlike mammalian RNase H1 (42).

**Figure 1.**
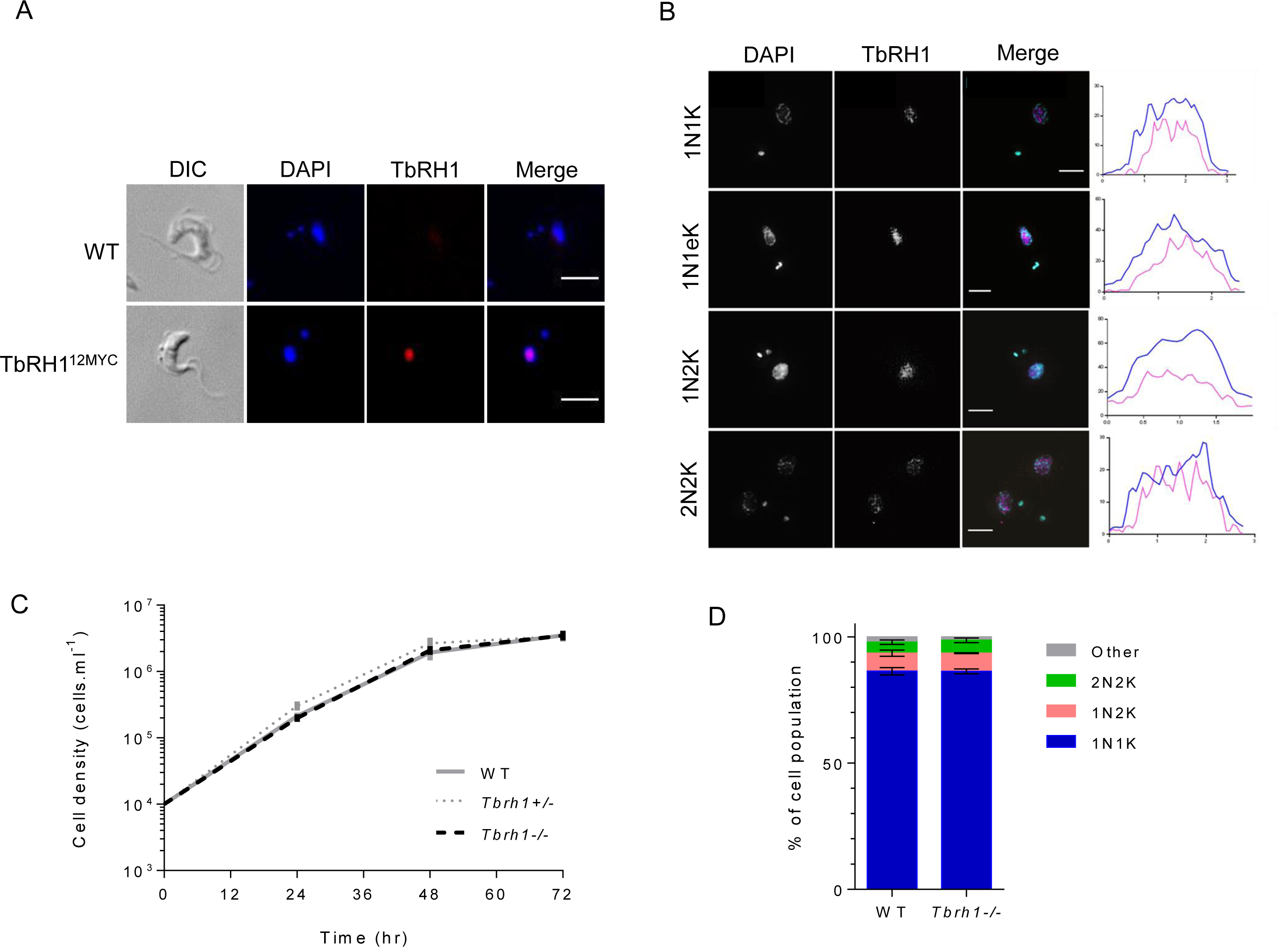
*T. brucei* ribonuclease H1 is a non-essential nuclear protein. A. Representative immunofluorescence images of *T. brucei* cells expressing Ribonuclease H1 (TbRH1) as a fusion with 12 copies of the myc epitope (TbRH1^12myc^); wildtype (WT) cells are shown for comparison. Anti-myc signal is shown in red and DNA is stained with DAPI (blue); merged DAPI and anti-myc signal is also shown, as is the cell outline (by differential interference contrast microscopy; DIC). Scale bars, 5 μm. **B.** Super-resolution structure-illumination imaging of TbRH1 and nuclear DNA colocalisation; TbRH1^12myc^ expressing cells are shown stained with anti-myc antiserum and with DAPI; representative images are shown of different cell cycles stages, and only in the merge of anti-myc (magenta) and DAPI (cyan) images is colour provided. Graphs plot length across the nucleus (x, pixels) versus mean pixel intensity at each position (y, arbitrary units) for DAPI (aqua) and TbRH1 (pink). Scale bars, 5 μm. **C.** Growth of WT cells and *T. brucei TbRH1* heterozygous (*TbRHl*+/-) or homozygous *Tbrh1*-/- mutants in culture, with mean population density shown at 24 hr intervals; error bars denote SD from three experiments. **D.** Percentage of the population of WT and *Tbrh1*-/- cells in discernible cell cycle stages, determined by DAPI staining and fluorescent imaging followed by counting the number and shape of nuclear (N) and kinetoplast (K) structures in individual cells: 1N1K, 1N2K and 2N2K and ‘other’ cells that do not conform to these patterns (>200 cells were counted for each cell type).

### R-loops accumulate throughout VSG transcription sites in the absence of RNase H1

To ask if TbRH1 targets R-loops within the VSG ES, we performed DNA-RNA immunoprecipitation coupled to next generation sequencing (DRIP-seq) (43) in both wild type (WT) and *Tbrh1*-/- cells, aligning DNA reads to the VSG ES using MapQ filtering (44) to ensure ES-specific mapping of Illumina short reads across regions of homology (Fig.2, Fig.S3). In WT cells there was limited read enrichment across the ES region spanning the promoter to the *VSG*, either in the actively transcribed ES (BES1, containing *VSG221*) or the 13 distinct silent ES. Pronounced enrichment was only observed proximal to the ends of the ES, downstream of the VSG, which most likely represents TERRA RNA since levels of the signal increased in *Tbrh1*-/- mutants (Figs.2A,E), the opposite of decreased TERRA RNA when TbRH1 is over-expressed (26). Loss of TbRH1 resulted in DRIP-seq signal throughout all ES, both active and silent (Fig.2A, Fig.S3). To check the mapping, we performed qPCR on DRIP samples (Fig.2B). Enrichment of sequence in the IP relative to input (non-IP) was substantially higher (^∼^10 fold) from *Tbrh1*-/- cells relative to WT for two *ESAGs* (6 and 8), confirming intra-ES R-loops. The same differential between mutant and WT was also seen with qPCR using primers recognising *VSG221* (BES1, active) or *VSG121* (BES3, inactive), confirming R-loops in both transcribed and untranscribed sites. Finally, on-bead treatment of samples with *E. coli* RNase HI prior to DNA recovery clearly reduced the IP enrichment in the *Tbrh1*-/- cells, confirming recovery of RNA-DNA hybrids.

**Figure 2.**
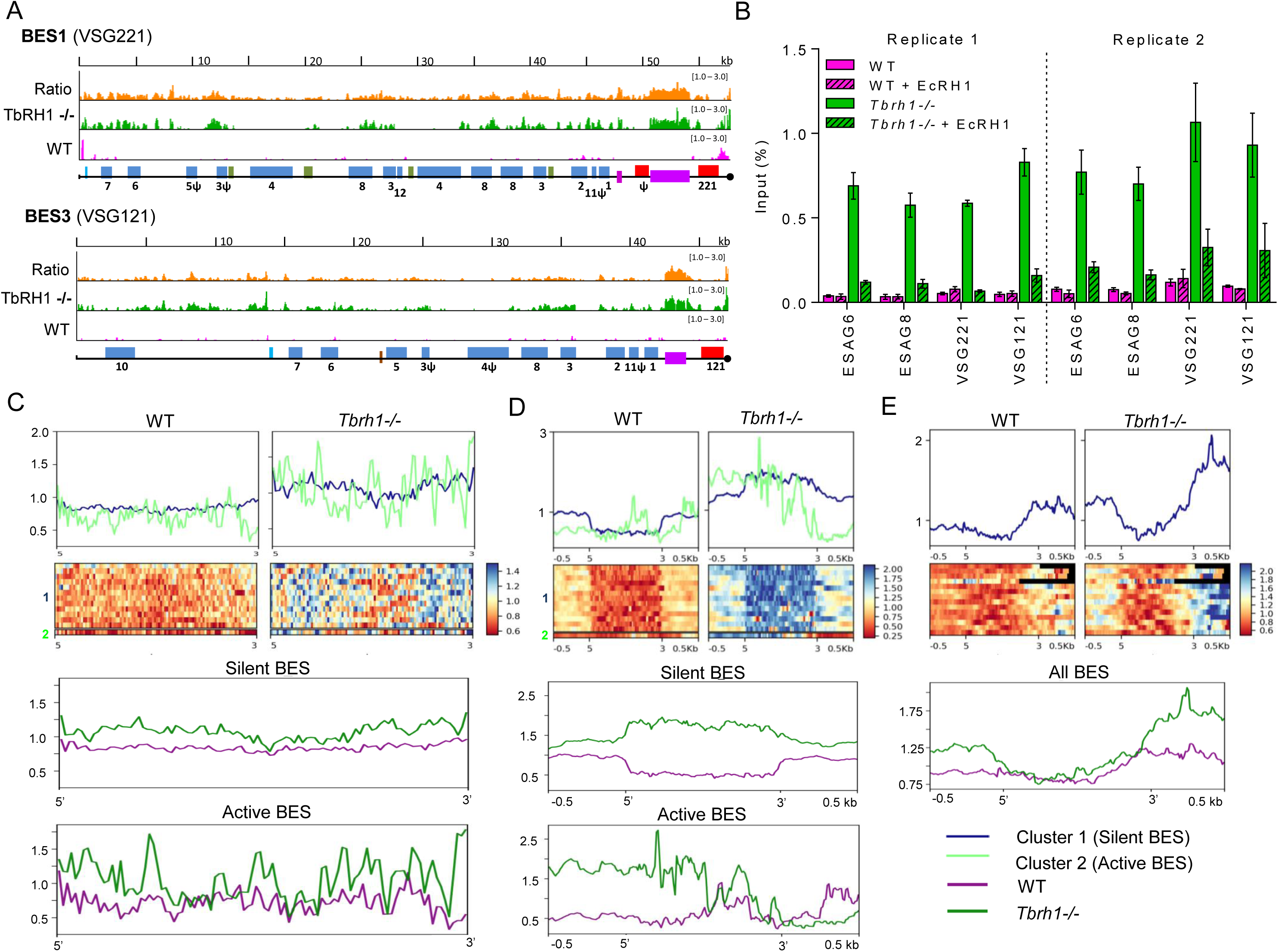
Localisation of R-loops in the Variant Surface Glycoprotein expression sites of *T. brucei* before after loss of Ribonuclease H1. A. Localisation of R-loops by DRIP-seq in wild type (WT) and *T. brucei* RNaseH1 homozygous mutant (*Tbrh1*-/-) bloodstream form cells. DRIP-seq signal is shown mapped to BES1 (the predominantly active VSG expression site (ES) of WT cells) and BES3 (which is mainly inactive). Pink and green tracks show normalised ratios of read-depth fold-change (1-3 fold) in IP samples relative to input in WT and *Tbrh1*-/- mutants, respectively, while the orange tracks show the ratio of IP enrichment in *Tbrh1*-/- cells compared with WT. Promoters (aqua), *ESAGs* (blue, numbered), 70-bp repeats (purple) and *VSGs* (red) are annotated as boxes; pseudogenes are indicated (ψ), hypothetical genes are shown in green, and the end of the available ES sequence is denoted by a black circle. **B**. DRIP-qPCR, with or without *E.coli* RNaseH1 (EcRH1) treatment, showing the percentage of PCR amplification in the IP sample relative to input for WT cells (pink) and *Tbrh1*-/- mutants (green); error bars display SEM for at least three technical replicates and data are shown for two biological replicates (1 and 2). **C-E**. DRIP-seq signal fold-change (IP relative to input samples; y-axes) plotted as heatmaps and average signal fold-change profiles over ES regions encompassing the *ESAGs* (C), 70-bp repeats (D) and *VSG* (E); for each region, 5’ and 3’ (x-axes) denote the upstream and downstream boundaries, and in some cases -/+ 0.5 kb of flanking sequence is shown. <u>Upper two panels</u>: comparison of WT and *Tbrh1*-/- DRIPseq signal using kmeans clustering, which separated the active (light green, cluster 2) and inactive (dark blue, cluster 1) ES when analysing *ESAGs* and 70 bp repeats, but not *VSGs.* <u>Lower panels</u>: Overlay of WT (purple) and *Tbrh1*-/- (green) DRIPseq signals in the three ES regions, with the active and silent ES displayed separately for the *ESAGs* and 70 bp repeats.

To examine the distribution and abundance of ES R-loops further, we performed K-means clustering of the read alignments from WT and *Tbrh1*-/- mutant DRIP-seq, separating the analysis into three ES components: the *ESAG*-containing region from the promoter to 70 bp repeats (Fig.2C), the 70 bp repeats (Fig.2D), and the *VSG* plus 500 bp of flanking sequence (Fig.2E). In all components of the ES, as expected, read abundance was greater in the *Tbrh1*-/- mutants than WT. However, the extent and pattern of enrichment was not equivalent in the three components, and nor was it always equivalent in active and silent ES. For the *ESAG* and 70 bp repeat components (Figs.2C, D), clustering analysis separated the active ES from all silent ES both in WT and *Tbrh1*-/- cells, suggesting differences dictated by transcription. In contrast, no such separation was seen around the *VSG*, where the active and silent ES could not be distinguished (Fig.2E). Indeed, the level of signal across the *VSG* ORFs was relatively low compared with upstream and, in particular, downstream (presumably telomeric) regions, despite low levels of R-loops being present (as confirmed by VSG DRIP-qPCR; Fig.2B). Enrichment of DRIP-seq signal in *Tbrh1*-/- cells relative to WT extended across the *ESAGs* (Fig.2C) and did not appear to be due to localisation to any specific sequence elements, such as the ORFs or untranslated intergenic regions (Fig.2A, Fig.S3), indicating R-loops became more abundant throughout the region of potential transcription upstream of the 70 bp in the absence of TbRH1. In contrast, the 70 bp repeats were notable for three features (Fig.2D). First, the level of enrichment across the repeats was notably higher in *Tbrh1*-/- mutants relative to WT than in both other components of the ES (^∼^2 fold in the 70 bp repeats, compared with ^∼^1.5 fold elsewhere; see also Fig.2A, Fig.S3). Second, in WT cells the small signal levels in the inactive ES were notably lower across the 70 bp repeats than surrounding sequence, whereas in *Tbrh1*-/- cells signal was greater across the repeats than the flanks. Third, a different pattern was seen in the active ES: in WT cells there was no DRIP-seq signal ‘dip’ within the 70 bp repeats, and in the *Tbrh1*-/- mutants the signal was more enriched at promoter-proximal repeats than telomere-proximal, following the direction of transcription. Taken together, the clustering analysis indicates TbRH1 plays a key role in removing R-loops in the VSG ES, with the 70 bp repeats being a focus for accumulation of the hybrids; furthermore, the distinct features of the mapping within the active ES relative to inactive ES suggest R-loop accumulation and removal by TbRH1 is co-transcriptional in WT cells.

### Loss of RNase H1 results in elevated levels of VSG coat switching

Given that R-loops accumulate within the VSG ES in the absence of TbRH1, we next asked if the increased abundance of the RNA-DNA hybrids is associated with altered VSG expression in *Tbrh1*-/- mutants relative to WT. To test this association, we first performed RT-qPCR to measure RNA levels of a selection of ES *VSGs* (Fig.3A). Five *VSGs* within silent ES in the Lister 427 *T. brucei* strain used here (14)(Fig.3A) displayed significantly higher RNA abundance in the *Tbrh1*-/- cells relative to WT. In addition, a small reduction (p value 0.054) in RNA levels was observed for *VSG221*, which is present in the predominantly active VSG ES in WT cells (BES1; Fig.2). To ask if RNA changes are limited to ES VSGs, we performed RNAseq on RNA from the WT and *Tbrh1*-/- cells and mapped the reads to all available annotated VSGs in the Lister 427 *T. brucei* strain (14, 16). In total, 63 VSGs displayed 1.5 fold or greater number of mapped RNAseq reads in the *Tbrh1*-/- mutants compared with WT (Fig.3B). Amongst these genes were nine bloodstream ES VSGs, of which four showed particularly pronounced read increases, though comparable changes in read depth were not obvious for the associated ESAGs within the ES (see VSG121 in BES3, Fig.S4). Increased RNAseq reads in the mutants were also detected for VSGs from all parts of the silent archive, with intact and pseudogenic array VSGs more frequently detected than metacyclic ES or minichromosomal VSGs (Figs. 3B,C). Taken together, the RT-qPCR and RNAseq data indicate that loss of RNaseH1 leads to increased levels of transcription from normally silent VSGs.

**Figure 3.**
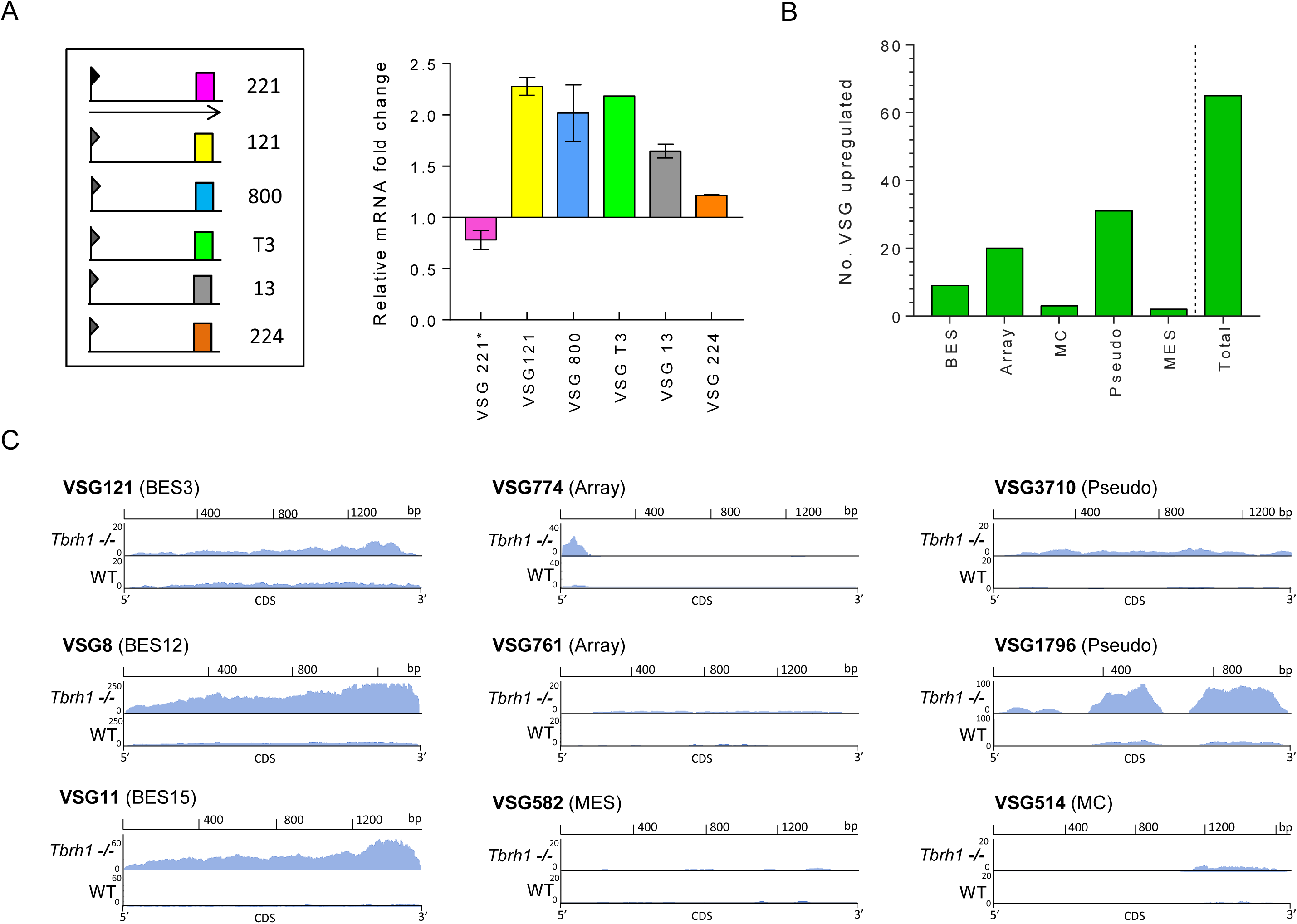
Loss of *T. brucei* RNase H1 results in increased transcription of silent VSGs. A. The left panel provides a simplified diagram of VSG expression sites (ES) used to generate a protective surface coat in the bloodstream *T. brucei* cells used in this study; only telomere-proximal *VSGs* (coloured boxes, numbered) from a selection of ES are shown, and the single ES being actively transcribed (encoding *VSG221*, pink) is denoted by an arrow extending from the promoter (flag). The right panel shows a graph of *VSG* RNA levels (corresponding to the ES diagram, and determined by RT-qPCR) in TbRH1 null mutants (*Tbrh1*-/- cells), plotted as fold-change relative to levels of the cognate *VSG* RNA in wild type cells (WT427 1.2); ^∗^ indicates *VSG221* is in the active ES of wild type cells, and error bars show SEM for three independent experiments. **B.** A graph depicting the number of VSG genes that display >1.5-fold increase in RNA abundance (determined by RNAseq, and normalised to gene length and total number of reads) in *Tbrh1*-/- cells relative to WT; the total number is sub-categorised depending on whether the VSGs have been localised to the bloodstream ES (BES), are intact genes in the subtelomeric arrays (array), are in mini-chromosomes (MC), are pseudogenes (pseudo), or are in the ES transcribed in the metacyclic life cycle stage (MES). **C.** Plots of normalised RNAseq read depth abundance (y-axes) relative to CDS position (x-axes) for a selection of the above VSGs; VSG identity numbers are from (16).

To test if these RNA changes extend to the VSG surface coat, we used immunofluorescence on unpermeabilised cells to evaluate the stability of VSG221 expression, since this VSG is normally resident in the predominantly transcribed ES (BES1). To do this, we first examined VSG221 expression over time, comparing the frequency with which three *Tbrh1*-/- and WT clones no longer expressed the protein during prolonged passage (Fig.4A). Despite the absence of immune selection against VSG221 expression, greater numbers of cells without surface VSG221 were seen in the *Tbrh1*-/- cells than in WT at each time point examined, indicating elevated levels of VSG switching throughout growth in culture. Notably, such elevated VSG switching is not associated with changes in population doubling time of the RNaseH1 mutants. To examine this effect in more detail, we next performed co-immunofluorescence (IF) on unpermeabilised WT and *Tbrh1*-/-cells (grown for >45 generations in culture) using antiserum recognising VSG221 (active BES1) or VSG121 (silent BES3). In this WT population, all cells analysed expressed VSG221 (Fig.3B), whereas ^∼^3.5% of the *Tbrh1*-/- mutant cells no longer expressed VSG221 on their surface (Fig.3B, C). Most of the cells (^∼^3.1%) that did not react with VSG221 antiserum also did not react with VSG121 antiserum, indicating they expressed a distinct VSG or VSGs on their surface. However, in a small proportion of cells (^∼^0.35%) VSG121 could be detectably expressed, indicating this gene had been activated. To determine if all VSG121-expressing *Tbrh1*-/- cells had switched off VSG221, we looked amongst the VSG121-expressing cells for costaining with both antisera (Fig.3C, D). As a control, immunofluorescence was also performed in a distinct WT strain (i.e. not lacking TbRH1) in which a transcription elongation blockade within BES1 has silenced this ES and predominantly activated BES3 (28, 45, 46), containing *VSG121* (Fig.3 C,D). Most (^∼^68%) *Tbrh1*-/- cells stained only with anti-VSG121 antiserum, though a minority (^∼^32%) were VSG221-VSG121 double expressers. Taken together, these data indicate that loss of TbRH1 results in an increased frequency at which expression of the active VSG is lost, which mainly reflects complete VSG switching events where the active VSG is no longer detected and expression of a distinct VSG occurs. Loss of mono-allelic control, which results in co-expression of VSGs from the active and at least one previously silent ES, is less common but was observed.

**Figure 4.**
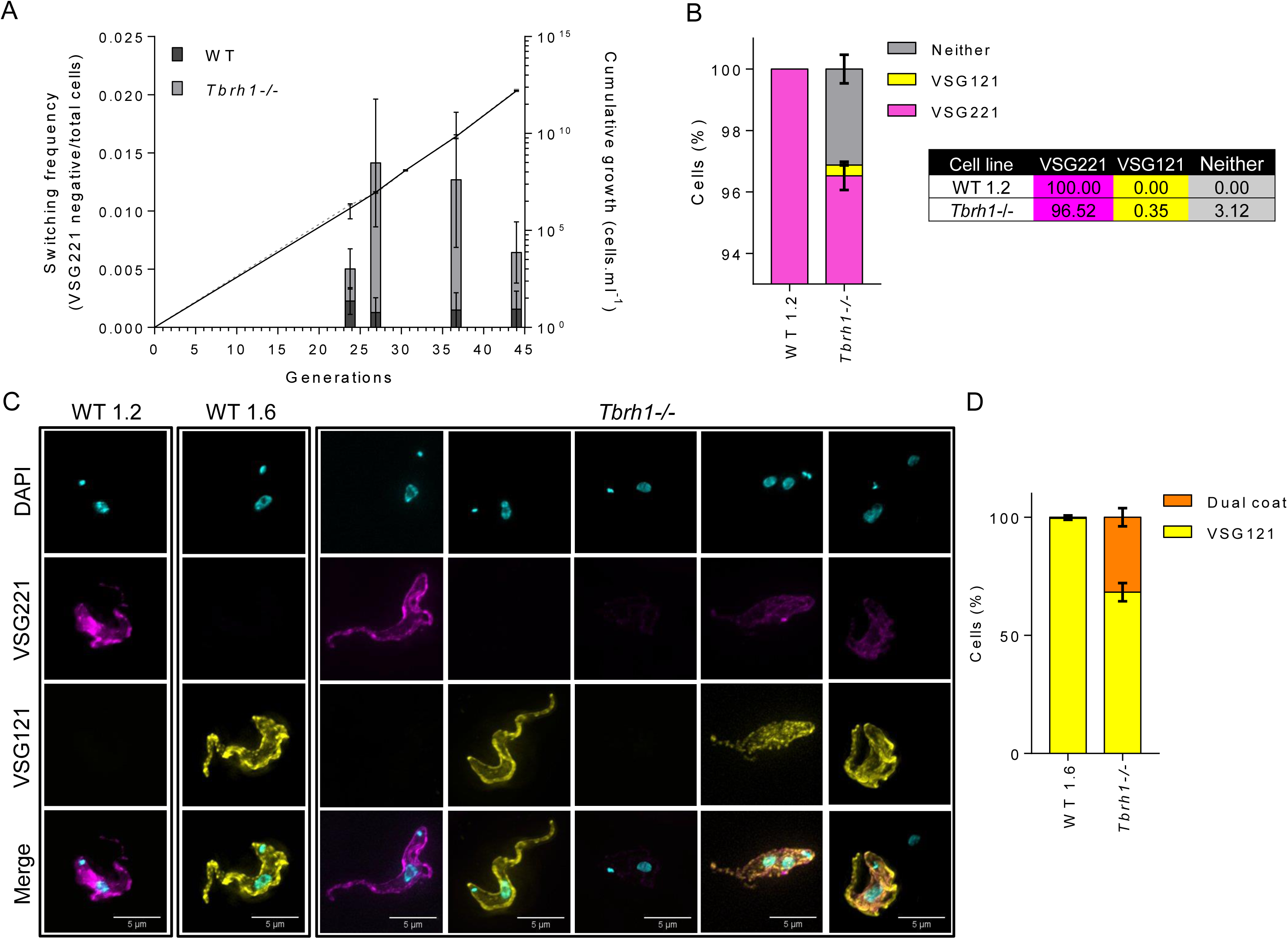
Loss of *T. brucei* RNase H1 induces VSG coat switching. A. Evaluation of the frequency at which *Tbrh1*-/- and WT cells switch off expression of VSG221 on their cell surface. For each cell type, three VSG221-expressing clones were generated and grown independently, by serial passage in culture, for a number of generations. At multiple time points the number of cells in the populations express or do not express VSG221 was assessed by immunofluorescence with anti-VSG221 antiserum; non-VSG221 expressing WT (black) and *Tbrh1*-/- (grey) cells are shown as a proportion of the total population (data shows average and SD of the three clones, and >200 cells were counted for each clone and at each time point). Cumulative density of WT (solid line) and *Tbrh1*-/- cells (dotted line) over the course of the analysis is shown; values depict average density and SD for the three clones of each cell type at the passages shown. **B** Percentage of WT (WT 1.2) and *Tbrh1*-/- cells expressing VSG221 or VSG121 on their surface, as determined by co-immunofluorescence imaging with anti-VSG221 and VSG121 antiserum. The graph depicts the relative proportions of cells in the population in which only VSG221 (magenta) or VSG121 (yellow) could be detected, as wells as cell with both (orange) or neither (grey) of the two VSG on their surface; >200 cells were analysed for each cell type in each of three replicates (error bars denote SEM). **C.** Co-immunofluorescence imaging of VSG221 and VSG121, providing examples of the surface coat configurations measured in (B); in addition to WT 1.2 cells and *Tbrh1*-/- mutants, an example of a cell is shown from a *T. brucei* strain (WT 1.6)(45) that predominantly expresses VSG121 and not VSG221 (Scale bars, 5 μm). **D.** Analysis of WT 1.6 and *Tbrh1*-/- mutant cells that express VSG121 on the cell surface, showing the percentages that simultaneously express VSG221 (orange) or only express VSG121 (yellow); >100 cells were analysed in each of three replicate experiments for each cell type.

### Loss of RNase H1 results in elevated levels of replication-associated genome damage

Two models might be considered to explain elevated VSG switching in *Tbrh1*-/- mutants relative to WT cells (Fig.5). In one model, ES R-loop accumulation impedes complete transcription of the active ES, selecting for cells in which a previously silent ES has been transcriptionally activated. Once activated, these newly expressed ES then accumulate R-loops, propagating the DRIP-seq signal across all ES (Fig.2). However, R-loops have also been linked to DNA breaks and rearrangement (39), through impeding DNA replication (47,51), as a result of elevating the levels of transcription-associated breaks (52-54), or because the RNA-DNA hybrids form in response to transcription-associated breaks (55-61). A second model, therefore, is that increased R-loops in *Tbrh1*-/- cells reflect the accumulation of damage in the ES, leading to recombination-based VSG switching. To try and separate these models, we compared levels of nuclear genome damage in WT and *Tbrh1*-/- cells by assessing expression of Thr130-phosphorylated histone H2A (γ-H2A), which increases in abundance after a range of genotoxic insults (62, 63) and in repair mutants (28). Western blotting revealed comparable levels of overall γ-H2A in WT and *Tbrh1*-/- cells (Fig.S5). However, IF analysis revealed a ^∼^2.3 fold increase in the number of *Tbrh1*-/- cells with detectable nuclear γ-H2A signal, rising from around 7% in WT (Fig.6A). Super-resolution structure-illumination microscopy revealed that in both WT and *Tbrh1*-/- cells most γ-H2A signal appeared as a single subnuclear focus, though some cells with >1 foci were present (Fig.6B; examples in Figs. 5D and S6). In addition, some cells displayed diffuse staining throughout the nucleus (Fig.6B), suggesting γ-H2A signal may represent various types of damage. DAPI staining of a *T. brucei* population provides a means to determine the cell cycle stage of individual cells, since replication and segregation of the nuclear (N) and kinetoplastid (K) genomes occur with different timings (64). In keeping with previous work (62) more WT cells displayed γ-H2A signal (Fig.6C) when they were undergoing nuclear replication (1N1eK) or were in G2-M phase (1N2K), with reduced numbers of signal-positive cells from the end of M phase (2N2K) through G1 (1N1K). Cell cycle quantification of the *Tbrh1*-/- mutants showed that the increased proportion of cells with γ-H2A signal was nearly entirely accounted for by greater numbers of 1N1eK (^∼^2.4-fold increase) or 1N2K (^∼^3.6-fold) cells with foci relative to WT (Fig.6C), indicating increased accumulation of nuclear damage occurs during replication of the genome in the absence of TbRH1.

**Figure 5.**
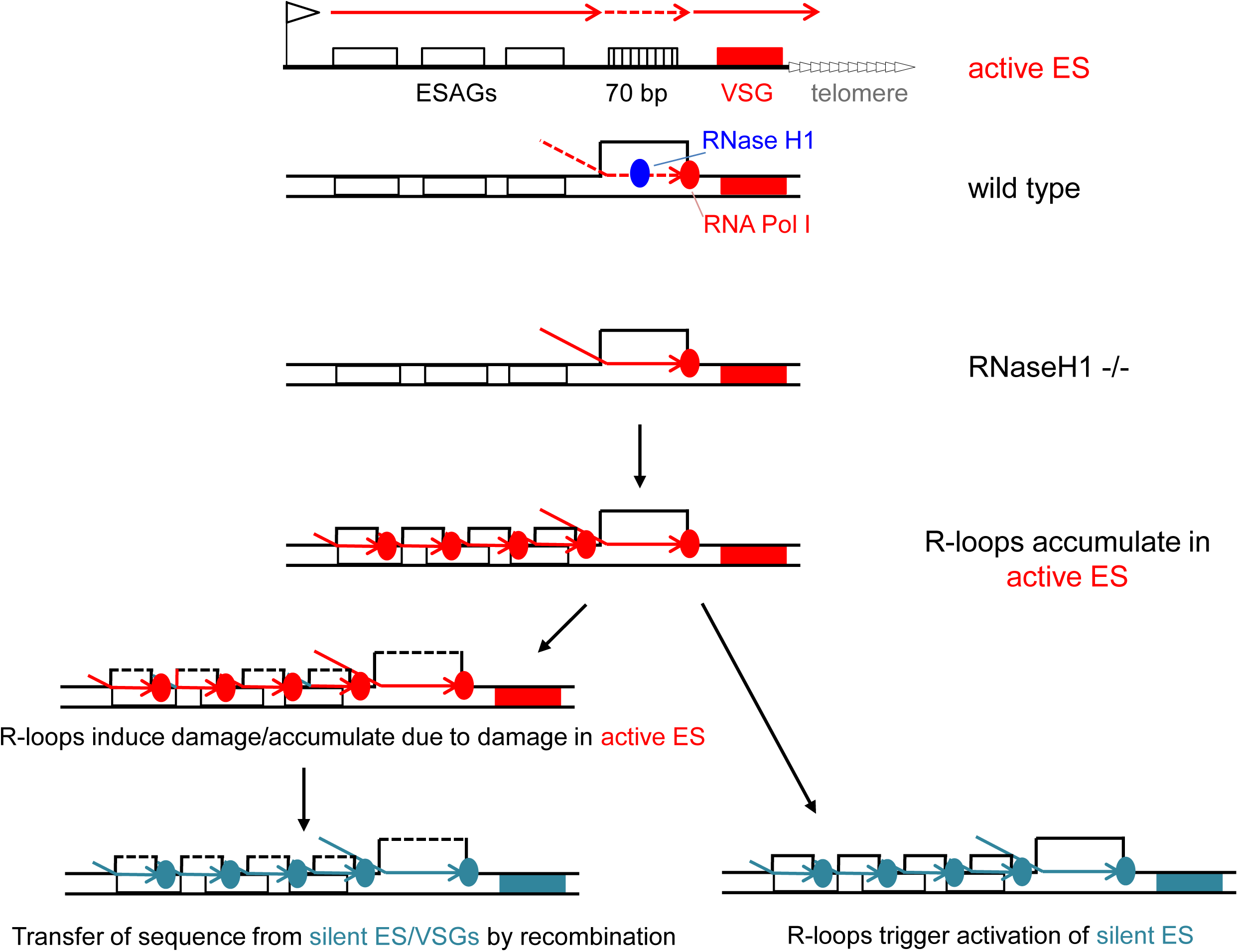
Model of R loop accumulation and resolution in the VSG expression sites of wild type and RNase H1 mutant *T. brucei* bloodstream form cells. The topmost diagram summarises the structure of and transcription across (red arrows) the active VSG expression site (ES, black line), including key features: telomere repeats (white arrows), the *VSG* gene (red box), 70 bp repeats (hatched box), *ESAG* genes (white boxes) and the promoter (flag). In the active ES transcription by RNA Polymerase (Pol) I must occur at high levels in order to generate sufficient VSG to form a dense surface coat. Pol I (red circle) passage is slowed when traversing the 70 bp repeats (dotted arrow), leading to the formation of R loops (DNA-RNA hybrid and extruded single strand DNA) that RNase H1 (blue circle) resolves to allow continued transcription. In TbRNaseH1 null mutants (-/-) R loops accumulate in the 70 bp repeats, obstructing transcription and leading to upstream RNA Pol I stalls and the formation of R loops across the length of the ES. Two outcomes of can then occur. In the first, reduced expression of the active ES and VSG is lethal, allowing selection for cells that have activated a distinct ES and VSG (green box). The lack of R-loop resolution by TbRNaseH1 in this site then results in R-loop accumulation and selection for activation of further ES (not shown). In the second outcome, R loops in the active ES lead to, or result from DNA damage, which is then repaired by homologous recombination, resulting in the transfer of sequence (e.g. *VSGs*, *ESAGs*) from the silent archive, including the silent ES, into the active ES.

**Figure 6.**
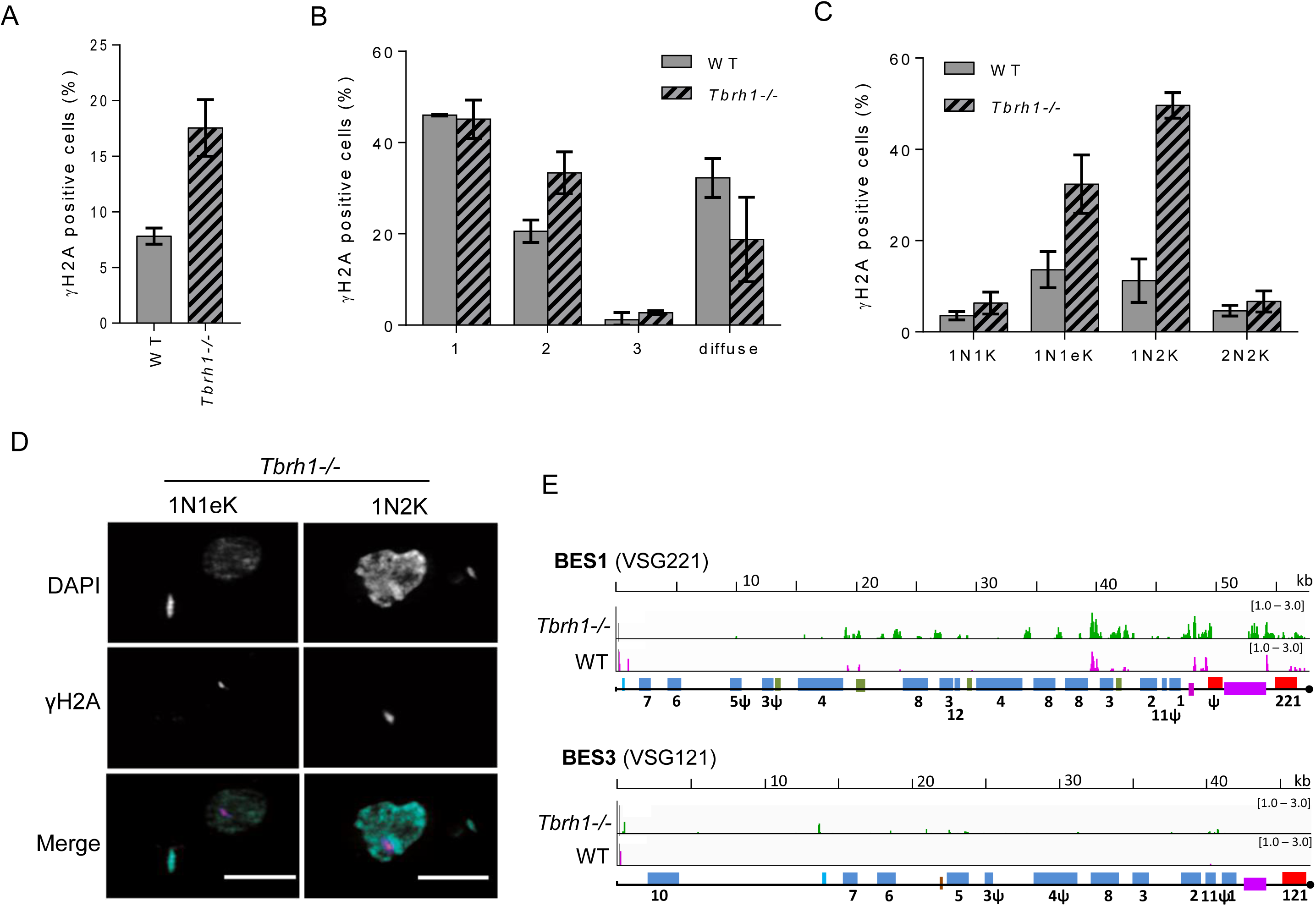
Loss of *T. brucei* RNase H1 leads to increased levels of nuclear damage in replicating cells and in the VSG expression site. A. Percentage of WT and *Tbrh1*-/- cells (n >200 in each of three replicates) that have detectable nuclear anti-γ-H2A signal. **B.** Distribution of intra-nuclear γ-H2A signal in WT and *Tbrh1*-/- cells (n >200 in each of three replicates). **C.** Percentage of cell cycle stages in WT and *Tbrh1*-/- cells with nuclear anti-γ-H2A signal; cell cycle stages were determined by DAPI staining and fluorescent imaging followed by counting the number and shape of nuclear (N) and kinetoplast (K) structures in individual cells: 1N1K, 1N1elongated K (1N1eK), 1N2K and 2N2K (n ≥50 for each cell cycle stage in three replicates)**. D.** Super-resolution structure-illumination microscopy imaging of anti-γ-H2A signal and co-localisation with DAPI in representative replicating (1N1eK and 1N2K; see below) *Tbrh1*-/- cells; only in the merge of anti-γ-H2A (magenta) and DAPI (cyan) images is colour provided (further examples of mutant and WT cells are provided in Fig.S5). **E.** Localisation of γ-H2A by ChiP-seq in WT and *Tbrh1*-/- cells. ChiP-seq signal is shown mapped to BES1 and BES3, which are represented as in Fig.2. Magenta and green tracks show normalised ratios of read-depth fold-change (1-3 fold) in IP samples relative to input in WT and *Tbrh1*-/- mutants, respectively.

### Loss of RNase H1 results in accumulation of DNA damage predominantly in the actively transcribed VSG expression site

To ask if some of the damage detected by microscopy of γ-H2A localises to the VSG ES, we performed ChIP-seq with anti-γ-H2A antiserum in WT and *Tbrh1*-/- cells, mapping the reads to the 14 ES using MapQ filtering (Fig.6E, Fig.S7). In WT cells low levels of γ-H2A ChIP reads were seen in all ES, but the level of enrichment was notably greatest and most widespread in the active ES (BES1, containing VSG221), in particular around the VSG and 70 bp repeat-proximal ESAGs. In the *Tbrh1*-/- cells, γ-H2A ChIP reads were detected at even greater levels in the active ES (BES1), both at the locations detected in WT cells and due to increased reads more proximal to the ES promoter. In contrast, though γ-H2A ChIP reads also increased in the silent ES of *Tbrh1*-/- cells (Fig.6E, Fig.S7), the extent of this change was more modest than was seen in the active ES. These data indicate DNA damage is present in the active ES, in particular proximal to the 70 bp repeats and *VSG*, where distinct assays have suggested the presence of DNA breaks (23-25). As the extent of γ-H2A signal increases after ablation of TbRH1, this response correlates with the increased abundance of R-loops, which may form preferentially in the active ES.

## Discussion

In this work we reveal that interplay between transcription and sequence composition of the VSG ES leads to R-loops acted upon by RNaseH1, and that loss of the ribonuclease results in increased replication-associated damage, including in the VSG ES, and increased VSG coat switching. These findings indicate that VSG ES structure lends itself to the generation of R-loops during the expression of the trypanosome’s crucial surface antigen, indicating RNA-DNA hybrids may be harnessed to provide pathogen-specific, discrete functions, such as antigenic variation.

The data we present here provide insight into the initiation of antigenic variation in *T. brucei* (Fig.5). We suggest that transcription through the VSG ES leads to the formation of R-loops that appear to be rapidly resolved, including by TbRH1, to ensure continued high rates of VSG coat expression (65). Within the ES the 70 bp repeats appear to be a pronounced site of VSG RNA-DNA hybrid accumulation, given the strongest R-loop ES enrichment is seen in this location in *Tbrh1*-/- mutants. R-loops may form more readily on the 70 bp repeats due to their sequence composition: individual 70 bp repeats show considerable size and sequence variation but are, in part, comprised of (TRR) repeats (15, 66) that can become non-H bonded (67) and promote recombination (68) when transcribed. Increased DRIP-seq signal in the active VSG ES of *Tbrh1*-/- mutants is therefore most readily explained by R-loops forming initially in the 70 bp repeats and extending back toward the RNA Pol I promoter due to retrograde spreading of the VSG ES transcription blockade. This scenario explains the less pronounced enrichment of R-loops within the telomere-proximal *VSGs.* In addition, it explains the lack of localisation of the R-loops to ES ORF or non-coding sequences, which contrasts with R-loop distribution in RNA Pol II and RNA Pol I multigene transcription units elsewhere in the genome, where R-loops show a very pronounced intragenic localisation (BIORXIV/2018/357020). Thus, it is very unlikely that R-loops in the ES are associated with transcription processing, but instead emerge from impaired RNA Pol I movement. Whether the R-loops form because transcription is impeded across the 70 bp repeats, or because of lesions within the repeats is unclear (see below).

Our data suggest that R-loop accumulation leads to VSG switching and spreading of R-loops into silent VSG ES by two routes: transcriptional and recombinational. The transcriptional model derives from the observation that RNAi-mediated loss of VSG expression from the active ES is lethal (65) and can select for switching (69). Co-transcriptional R-loop formation might itself lead to reduced transcription of the active VSG ES, or R-loops might form because of lesions generated during ES transcription. Irrespective, once R-loops form in the ES, in particular in the absence of TbRH1, they would provide a blockade to full transcription, causing further R-loops to form in the ES and providing a selection for activation of a new ES. Once a new ES is activated, transcription in this locus then gradually suffers the same R-loop blockade, which is exaggerated by the absence of TbRH1. The model is consistent with the presence of cells expressing both VSG221 and VSG121, since co-expression of at least two VSGs indicates altered transcriptional control. However, such double expressers are relatively rare and the seemingly equivalent accumulation of R-loops in both the active ES and in all silent ES is not simple to reconcile with the continued predominant transcription of VSG221 (BES1) in the *Tbrh1*-/- mutants. Therefore a second possibility is that R-loops do not merely reflect or cause transcription blockade, but are associated with recombination of silent VSGs, and in some circumstances *ESAGs* from silent ES, into the active ES. An R-loop VSG recombination model is consistent with the greater levels of γ-H2A signal in the active ES and, in this scenario, the relatively similar level of R-loop enrichment in both the active and silent ES could be explained by gene conversion of silent ES sequences into the active ES. However, the lack of evidence for increased RNA spanning the silent ES (such as BES3) in the *Tbrh1*-/- mutants contrasts with increased RNAseq reads for multiple ES VSGs, and does not readily match the greater density of γ-H2A signal proximal to the 70 bp repeats in the mutants. Instead, the γ-H2A ChIP mapping appears consistent with damage initially arising in the telomere-proximal region of the ES, perhaps due the nature of the repeats (above). If correct, this may explain the RNAseq data, with telomere-proximal damage leading to activation not simply (or even predominantly) of silent ES VSGs, but any VSG in the silent archive. In this model, accumulation of R-loops throughout the silent ES may indicate *trans* formation of the RNA-DNA hybrids after their generation in the active site, rather than transcription-associated formation. Irrespective of the precise details, the accumulated data presented here suggests that R-loop-associated recombinational switching predominates over R-loop-associated transcriptional switching. Nonetheless, since we could detect *Tbrh1*-/- cells in which at least two VSGs were expressed on the cell surface, a commonality of R-loops acting during transcription-mediated and recombination-mediated antigenic variation in *T. brucei* is possible. In other words, these reactions may not be entirely independent in mechanisms but, instead, share a common initiating event. Such commonality might explain why ablation of the homologous recombination factors RAD51, BRCA2 and RAD51-3 impairs both VSG gene conversion and transcriptional switching (70-72), as well as why induction of DNA damage can elevate levels of silent VSG expression (73).

Precisely how R-loops intersect with recombinational VSG switching will need to be explored further, but several routes might be considered, which have intriguing parallels in other eukaryotes. Accumulation of R-loops due to ES transcription pausing may alone be enough to generate the increased damage we detect in the active ES (and thereby induce recombination), perhaps due to prolonged negative supercoiling downstream of RNA Pol I increasing the likelihood of R-loop formation and greater exposure of the single-stranded DNA in the RNA-DNA hybrid. Alternatively, R-loops may not be the cause of damage but may form in response to the formation of lesions, perhaps due to ES transcription, detected by γ-H2A. Both routes may explain detection of putative DSBs within VSG ES (23-25). Determining whether R-loops cause or respond to DNA breaks remains challenging in any setting (38, 55), since it is clear that chromatin and repair pathways can modulate the damaging effects of R-loops (74-77), while at the same time single- or double-stranded DNA breaks can induce R-loop formation (56, 61). A final route that might be considered is that R-loops do not merely affect transcription, but impede DNA replication through the active VSG ES and lead to breaks that elicit a VSG switch. This final route would be compatible with the elevated levels of γ-H2A in replicating *Tbrh1*-/- cells, and with previous observations showing the active VSG ES replicates earlier than all the silent ES (28). In other words, it is possible that the ES structure and function has evolved to target replication-transcription clashes to the site of VSG expression to facilitate switching. Currently none of these scenarios can be ruled out, but the effects of TbRH1 loss on inducing VSG switching appears consistent with observations in other eukaryotes, and may explain the previously characterised roles of several repair factors. Mutation of *T. brucei* RAD51, the key catalytic enzyme of homologous recombination, impairs VSG switching (70). Intriguingly, yeast Rad51 has recently been shown to promote formation of R-loops and rearrangement at certain loci (78), an effect that is abrogated when factors that promote Rad51 activity are mutated, consistent with the impairment of VSG switching in *T. brucei* BRCA2 and RAD51-3 mutants (71, 72). In both yeast and mammals, loss of the RecQ helicases Sgs2 and BLM, respectively, causes elevated levels of R-loops and locus-specific instability (79), a response that may explain increased VSG switching by gene conversion when RECQ2 (the *T. brucei* orthologue) is mutated (28). Moreover, the R-loop role of Sgs2/BLM has been interpreted as being necessary to tackle replication-transcription clashes, a role that may explain the distinct phenotypes of *T. brucei* RECQ2 mutants when acting on DNA double strand breaks and during VSG switching (28). Finally, mutation of both RNase H enzymes in yeast has been documented to cause elevated levels of DNA damage (detected as RAD52 localisation) at rRNA genes, an effect that is due to RNA Pol I transit and results in gene conversion by break-induced replication, a process that has been suggested to mediate VSG switching in the RNA Pol I-transcribed ES (80, 81). Despite the broad parallels between these observations and the demonstration of RNA-DNA hybrids within the VSG ES, it is probably premature to draw clear mechanistic parallels. For instance, it has long been known that R-loops mediate mammalian immunoglobulin gene class switching (82), but the nature of transcription and lesion generation (7) do not have obvious similarities with *T. brucei* VSG switching. To understand the relationship between R-loops and VSG switching, a number of questions need to be addressed, including: do R-loops generate or follow from ES lesions; what form of DNA lesion results from, or generates the R-loops; how are R-loop-associated lesions signalled to initiate repair; and do increased levels of R-loops in the silent ES indicate movement of the hybrids *in trans* from the active ES? Irrespective of the detailed mechanism, positioning of the 70 bp repeats immediately upstream of the *VSG* appears advantageous, targeting breaks to allow recombination-mediated break repair to access any of the ^∼^1000 *VSGs* outside the VSG ES. R-loops that form in other locations in the active VSG ES would drive intra-ES recombination, which is observed frequently (83).

Beyond the proposed mechanistic involvement of R-loops in directing VSG switching, the data in this work reveal wider overlap with emerging roles for RNA in many immune evasion strategies. Antigenic variation in *Neisseria gonorrhoeae* relies upon the expression of a small non-coding RNA, upstream of and antisense to the *pilE* expression site, across a guanine quartet-forming DNA sequence (84), which results in DNA nicks that may elicit recombination (85). Intriguingly, small RNAs may also be generated from silent *pilS* recombination substrates (86). Non-coding RNA is widespread in *Plasmodium*, including sense and antisense non-coding RNA (ncRNA) that emanates from the promoter (87) and intron (88) of *var* genes, which mediate antigenic variation, as well as from a *var*-associated GC-rich ncRNA gene family (89). Modulation of the expression of the ncRNAs, as well as mutating a novel exoribonuclease (87), undermines the transcriptional controls that determine singular *var* gene expression during antigenic variation. Though it has not to date been reported that R-loops form in these settings, as we describe for *T. brucei*, and the generation and action of effector transcripts is very likely to be organism-specific, it is notable that both recombination and transcription events during antigenic variation are influenced by RNA, which in at least two cases interacts with DNA. Characterising the factors and reactions that act on the *T. brucei* ES R-loops to dictate the dynamics of antigenic variation will reveal how similar or distinct the processes are in the different pathogens.

## Methods

### Molecular and cell biology techniques

All cell lines used were bloodstream form parasites, which were maintained in HMI-9 medium supplemented with 10% (v/v) FBS (Sigma-Aldrich, Missouri, USA) and 1% (v/v) of penicillin-streptomycin solution (Gibco) at 37°C and 5% CO2. C-terminus endogenous tagging of TbRH1 was carried out as previous described (90). Briefly, the C-terminal 626 bp sequence of the *TbRH1* ORF was PCR-amplified using primers CGACG*AAGCTT*CTGCGGATGACGGTAATG and CGACG*AGATCT*TGTGAATCGCCCTTTGGC and cloned into the pNATx12M plasmid containing 12 copies of the c-myc epitope. The construct was then stably transfected into *T. brucei brucei* Lister 427 MITat1.2 cells after digestion with *PstI*. Heterozygous (-/+) and homozygous (-/-) *Tbrh1* knockout cell lines were generated using two constructs containing cassettes of either blasticidin or neomycin resistance genes between α-β tubulin and actin intergenic regions, flanked by sequences homologous to the 5’ and 3’ UTRs of *TbRH1*, essentially as described in (28). Homologous flanking regions were PCR-amplified using the following primers: 5’ UTR CGACG*GGATCC*TTGCCTTACCCGTGTTTT and CGACG*TCTAGA*CCTTTTCTTTCCCATGGAC, 3’ UTR CGACG*CCCGGG*AGGTGTGTATGGGAATGA and CGACG*CTCGAGG*CACCACCCAGTATAGAAA. Total RNA was extracted using an RNeasy Mini Kit (Qiagen) and reverse transcribed with SuperScript II Reverse Transcriptase (Invitrogen) using random hexamer primers. Power SYBR Green Master Mix (Invitrogen) was used to perform qPCR and fold change was calculated using the 2-ΔΔCT method (91).

### ChIP-seq analysis

Both DRIP and γH2A ChIP sample preparation was performed using a ChIP-IT Enzymatic Express kit (Active Motif). Briefly, ^∼^ 2×10^8^ cells were grown to log phase before fixing in 1% formaldehyde for 5 min whilst shaking at room temperature, before 1 mL of 10X Glycine Buffer was added directly to the cells to stop fixation. Cells were then pelleted, re-suspended in Glycine Stop-Fix Solution and shaken at room temperature for 5 min. Cells were next lysed, according to the manufacturer’s protocol, allowing chromatin to be extracted and digested for 5 min with Enzymatic Shearing Cocktail at 37 °C to produce ^∼^200 bp fragments. IP was performed overnight at 4 °C with 4.5 ng of S9.6 (Kerafast) or 3 μg of anti-γH2A antibody. For DRIP, on-bead treatment of control DRIP samples was performed as previously described (43). qPCR was performed directly from DNA recovered from DRIP samples, with or without EcRH1 treatment, using SYBR Select Master Mix (Invitrogen). The amount of DNA in IP samples was expressed as a percentage of input DNA, using CT values first adjusted by the dilution factor of each sample.

Library preparation was performed using a TruSeq ChIP Library Preparation Kit (Illumina) and fragments of 300 bp, including adaptors, were selected with Agencourt AMPure XP (Beckman Coulter). Sequencing was performed with an Illumina NextSeq 500 platform. Reads were trimmed using TrimGalore (https://github.com/FelixKrueger/TrimGalore) under default settings before alignment to the Lister 427 bloodstream VSG expression sites using Bowtie2 (92) in “very-sensitive” mode. Reads with a MapQ value <1 were removed using SAMtools (93), leaving at least 30 million aligned reads per sample. The fold change between input and IP read depth was determined for each sample using the DeepTools bamCompare tool (library size was normalised by read count and fold change was expressed as a ratio) and visualised as tracks with IGV (94). Normalised ratio files were also used to generate plots and perform kmeans cluster analysis using deepTools computeMatrix, plotProfile and plotHeatmap (95) tools.

### RNA-seq analysis

For RNA -seq analysis, total RNA was extracted using the RNeasy Mini Kit (Qiagen). Poly(A) selection and library preparation was then performed using the TruSeq Stranded Total RNA kit (Illumina) and sequencing of 75 bp paired-end reads was performed using the Illumina NextSeq 500 platform.

RKPM was calculated for each available VSG coding region (16) ignoring duplicate reads. Fold-change in RPKM for each VSG was calculated for *Tbrh1*-/- relative to WT. To ask what transcripts displayed altered levels in the *Tbrh1*-/- mutants relative to WT cells fold in RPKM was determined in RStudio. RNAseq reads were aligned to the Lister 427 VSG ES (14) and annotated VSGs (16) using HISAT2 (96) in ‘no splice alignment’ mode; reads with a MapQ value <1 were removed using SAMtools, which has been shown to remove >99% of short read alignment to the wrong ES (44). Read mapping was visualised using Matplotlib and a custom python script.

### Immunofluorescence

VSG immunofluoresence analysis was performed as previously described (18). Briefly, cell were fixed in 1% formaldehyde (FA) at room temp for 15 min. Cells were then blocked in 50% foetal bovine serum (FBS) for 15 min before primary (α-VSG221, 1:10000; α-VSG121, 1:10000: gift from D. Horn) and secondary (Alexa Fluor 594 goat α-rabbit (Molecular Probes) 1:1000; Alexa Fluor 488 goat anti-rat (Molecular Probes), 1:1000) antibody staining was carried out at room temp for 45 mins in both cases. Cells were then mounted in Fluoromount G with DAPI (Cambridge Bioscience, Southern Biotech). For 12myc-TbRH1 and yH2A staining, cells were first adhered to slides before fixing in 4% FA for 4 min and then quenched in 100 nM glycine. Cells were permeabilised in 0.2% triton-X 100 for 10 min. Blocking was performed for 1 hour with 3% FBS before antibody staining (α-myc Alexa Fluor 488 conjugated (Millipore), 1:500; α-γH2A, 1:1000, and Alexa Fluor 488 goat α-rabbit (Molecular Probes), 1:1000) and mounting in DAPI as described above. For counting purposes cells were imaged using an Axioscope 2 fluorescence microscope (Zeiss) with a 60x objective. Higher resolution of VSG staining was performed with a DeltaVision Core Microscope (Applied Precision), using a 100x 1.4 oil objective (Olympus). Super-resolution structured-illumination imaging of 12myc-RH1 and yH2A signal was performed using an Elyra PS.1 microscope (Carl Zeiss) using a 63x 1.4 objective.

## Acknowledgements

We thank all lab members for invaluable discussions, and David Horn and Sebastian Hutchinson for comparing unpublished data.

## Data Access

Sequences used in the mapping have been deposited in the European Nucleotide Archive (accession number PRJEB21868). DRIPseq analysis will be hosted at TriTryDB (http://tritrypdb.org/tritrypdb/) in an upcoming release.

## Supporting Information

**Figure S1. Ribonuclease H1 is a nuclear protein in bloodstream form *T. brucei* parasites. A.** Western blot analysis, using anti-myc antiserum, of a *T. brucei* clonal cell expressing C-terminally 12myc epitope-tagged TbRH1 from the endogenous locus in bloodstream form *T. brucei;* untagged, wild type (WT) cells are shown for comparison, and the estimated size of TbRH1-12myc is indicated (kDa). **B.** Fluorescence signal intensity (a.u., arbitrary units) of DAPI (cyan) and anti-myc signal (magenta) in TbRH1-12myc expressing cells, separated into different discernible cell cycle stages (determined by number and shape of nuclear (N) and kinetoplast (K) structures seen after DAPI staining: 1N1K, 1N1elongatedK (1N1eK), 1N2K and 2N2K); dots denote intensity of individual cells and the median values (horizontal lines) interquartile range (error bars) are shown. Significance was determined by Kruskal-Wallis non-parametric test: (^∗^) p-value <0.05; (^∗∗^) p-value < 0.01; (^∗∗∗^) p-value < 0.001; (^∗∗∗∗^) p-value <0.0001.

**Figure S2.Generation and validation of *TbRH1* null mutants in bloodstream form *T. brucei.* A.** confirmation of replacement of the *TbRH1* open reading frame (ORF) with selective neomycin (*NEO*) and blasticidin (*BSD*) resistance gene cassettes. The upper two gels show PCR targeting the 3’ end of *NEO* or *BSD* resistance genes, testing linkage of these genes to *TbRH1* flanks after cassette insertion into the *TbRH1* locus. The lowest gel shows PCR of part of the *TbRH1* ORF in wild type cells (WT427), in a *NEO* transformant (Tbrh1+/- neo) and in two *NEO* and *BSD* transformants (Tbrh1 null CL1 and CL2). **B.** RT-qPCR of TbRH1 RNA levels, comparing abundance in WT427, *Tbrh1+/-* cells and in *Tbrh1*-/- null mutants; RNA levels in WT cells (relative to a control RNA) were set at 100% and levels in the mutants are shown as a percentage (error bars show SD from three experiments).

**Figure S3. DRIP-seq analysis of all bloodstream VSG expression sites characterised in *T. brucei* strain Lister 427.** DRIP-seq was performed with wild type (WT) and *Tbrh1*-/- cells and reads mapped to all ES (BES, numbered as in (14)) not shown in Fig.1. Promoters (aqua), *ESAGs* (blue), 70-bp repeats (purple), *VSGs* (red), non-*ESAG* genes (green) and a drug resistance gene (navy) are annotated as boxes. Pink and green tracks show normalised ratios of read-depth enrichment in IP samples relative to input in WT and *Tbrh1*-/- mutants, respectively, while the orange tracks show the ratio of IP enrichment in *Tbrh1*-/- cells compared with WT.

**Figure S4. Bloodstream expression site RNA abundance in the presence and absence of RNaseH1.** RNAseq read depth is shown in WT cells and *Tbrh1*-/- mutants across the length of two VSG ES (BES1 and BES2, containing VSG221 and VSG121, annotated as in Fig2A); note the read depth scale (y-axes) for each is distinct for the ES regions containing the VSG (red box) and the ESAGs (blue boxes, numbered). For all other ES, RNAseq read depth (normalised to gene length and total number of reads) is shown only for the VSGs, which are numbered according to (16), with the ES that houses them indicated (see Fig.S3).

**Figure S5. Expression levels of γ-H2A in *T. brucei* RNasaeH1 mutants do not change substantially compared with wild type cells. A.** Western blot of γ-H2A, detected by specific antiserum, in wild type (WT) and *T. brucei* RNaseH1 null mutants (*Tbrh1*-/-); antiserum detecting EF1-α provides a loading control. **B.** Relative density of γ-H2A western blot signal, normalised to EF1-α, is compared in WT (normalised to 1.0) and *Tbrh1*-/- cells.

**Figure S6. Subnuclear γ-H2A foci in replicating *T. brucei* RNasaeH1 mutants and wild type cells.** Super-resolution structure-illumination immunofluorescent imaging of anti-γ-H2A signal and co-localisation with DAPI in a number of replicating (1N1eK and 1N2K) *Tbrh1*-/- and wild type (WT) *T. brucei* bloodstream cells; only in the merge of anti-γ-H2A (magenta) and DAPI (cyan) images is colour provided. Scale bars, 5 μm.

**Figure S7. γ-H2A localisation in all bloodstream VSG expression sites characterised in *T. brucei* strain Lister 427.** ChIP-seq was performed with specific antiserum against γ-H2Ain wild type (WT) and *Tbrh1*-/- cells and Illumina reads mapped to all ES (BES, numbered as in(14)) not shown in Fig.1. Promoters (aqua), *ESAGs* (blue), 70-bp repeats (purple), *VSGs* (red), non-*ESAG* genes (green) and a drug resistance gene (navy) are annotated as boxes. Pink and green tracks show normalised ratios of read-depth enrichment in IP samples relative to input in WT and *Tbrh1*-/- mutants, respectively.

## References

1. Yao MC, Chao JL, Cheng CY. Programmed Genome Rearrangements in Tetrahymena. Microbiology spectrum. 2014;2(6).

2. Sterkers Y, Crobu L, Lachaud L, Pages M, Bastien P. Parasexuality and mosaic aneuploidy in Leishmania: alternative genetics. Trends in parasitology. 2014;30(9):429–35.

3. Ubeda JM, Raymond F, Mukherjee A, Plourde M, Gingras H, Roy G, et al. Genome-wide stochastic adaptive DNA amplification at direct and inverted DNA repeats in the parasite Leishmania. PLoS biology. 2014;12(5):e1001868.

4. Lee CS, Haber JE. Mating-type Gene Switching in Saccharomyces cerevisiae. Microbiology spectrum. 2015;3(2):MDNA3-0013-2014.

5. Klar AJ, Ishikawa K, Moore S. A Unique DNA Recombination Mechanism of the Mating/Cell-type Switching of Fission Yeasts: a Review. Microbiology spectrum. 2014;2(5).

6. Roth DB. V(D)J Recombination: Mechanism, Errors, and Fidelity. Microbiology spectrum. 2014;2(6).

7. Hwang JK, Alt FW, Yeap LS. Related Mechanisms of Antibody Somatic Hypermutation and Class Switch Recombination. Microbiology spectrum. 2015;3(1):MDNA3-0037-2014.

8. Deitsch KW, Lukehart SA, Stringer JR. Common strategies for antigenic variation by bacterial, fungal and protozoan pathogens. NatRevMicrobiol. 2009;7(7):493–503.

9. Palmer GH, Brayton KA. Gene conversion is a convergent strategy for pathogen antigenic variation. Trends Parasitol. 2007;23(9):408–13.

10. Vink C, Rudenko G, Seifert HS. Microbial antigenic variation mediated by homologous DNA recombination. FEMS microbiology reviews. 2012;36(5):917–48.

11. Obergfell KP, Seifert HS. Mobile DNA in the Pathogenic Neisseria. Microbiology spectrum. 2015;3(1):MDNA3-0015-2014.

12. Gunzl A, Kirkham JK, Nguyen TN, Badjatia N, Park SH. Mono-allelic VSG expression by RNA polymerase I in Trypanosoma brucei: expression site control from both ends? Gene. 2015;556(1):68–73.

13. Glover L, Hutchinson S, Alsford S, McCulloch R, Field MC, Horn D. Antigenic variation in African trypanosomes: the importance of chromosomal and nuclear context in VSG expression control. Cellular microbiology. 2013;15(12):1984–93.

14. Hertz-Fowler C, Figueiredo LM, Quail MA, Becker M, Jackson A, Bason N, et al. Telomeric expression sites are highly conserved in Trypanosoma brucei. PLoS ONE. 2008;3(10):e3527.

15. Hovel-Miner G, Mugnier MR, Goldwater B, Cross GA, Papavasiliou FN. A Conserved DNA Repeat Promotes Selection of a Diverse Repertoire of Trypanosoma brucei Surface Antigens from the Genomic Archive. PLoS genetics. 2016;12(5):e1005994.

16. Cross GA, Kim HS, Wickstead B. Capturing the variant surface glycoprotein repertoire (the VSGnome) of Trypanosoma brucei Lister 427. Molecular and biochemical parasitology. 2014;195(1):59–73.

17. Marcello L, Barry JD. Analysis of the VSG gene silent archive in Trypanosoma brucei reveals that mosaic gene expression is prominent in antigenic variation and is favored by archive substructure. Genome Res. 2007;17(9):1344–52.

18. Glover L, Hutchinson S, Alsford S, Horn D. VEX1 controls the allelic exclusion required for antigenic variation in trypanosomes. Proceedings of the National Academy of Sciences of the United States of America. 2016;113(26):7225–30.

19. Batram C, Jones NG, Janzen CJ, Markert SM, Engstler M. Expression site attenuation mechanistically links antigenic variation and development in Trypanosoma brucei. eLife. 2014;3:e02324.

20. Figueiredo LM, Janzen CJ, Cross GA. A histone methyltransferase modulates antigenic variation in African trypanosomes. PLoS Biol. 2008;6(7):e161.

21. Cestari I, Stuart K. Inositol phosphate pathway controls transcription of telomeric expression sites in trypanosomes. Proceedings of the National Academy of Sciences of the United States of America. 2015;112(21):E2803–12.

22. McCulloch R, Morrison LJ, Hall JP. DNA Recombination Strategies During Antigenic Variation in the African Trypanosome. Microbiology spectrum. 2015;3(2):MDNA3-0016-2014.

23. Boothroyd CE, Dreesen O, Leonova T, Ly KI, Figueiredo LM, Cross GA, et al. A yeast-endonuclease-generated DNA break induces antigenic switching in Trypanosoma brucei. Nature. 2009;459(7244):278–81.

24. Glover L, Alsford S, Horn D. DNA break site at fragile subtelomeres determines probability and mechanism of antigenic variation in African trypanosomes. PLoS pathogens. 2013;9(3):e1003260.

25. Jehi SE, Wu F, Li B. Trypanosoma brucei TIF2 suppresses VSG switching by maintaining subtelomere integrity. Cell research. 2014;24(7):870–85.

26. Nanavaty V, Sandhu R, Jehi SE, Pandya UM, Li B. Trypanosoma brucei RAP1 maintains telomere and subtelomere integrity by suppressing TERRA and telomeric RNA:DNA hybrids. Nucleic Acids Res. 2017.

27. Hovel-Miner GA, Boothroyd CE, Mugnier M, Dreesen O, Cross GA, Papavasiliou FN. Telomere Length Affects the Frequency and Mechanism of Antigenic Variation in Trypanosoma brucei. PLoS Pathog. 2012;8(8):e1002900.

28. Devlin R, Marques CA, Paape D, Prorocic M, Zurita-Leal AC, Campbell SJ, et al. Mapping replication dynamics in Trypanosoma brucei reveals a link with telomere transcription and antigenic variation. eLife. 2016;5.

29. Devlin R, Marques CA, McCulloch R. Does DNA replication direct locus-specific recombination during host immune evasion by antigenic variation in the African trypanosome? Current genetics. 2016.

30. Santos-Pereira JM, Aguilera A. R loops: new modulators of genome dynamics and function. Nature reviews Genetics. 2015;16(10):583–97.

31. Stuckey R, Garcia-Rodriguez N, Aguilera A, Wellinger RE. Role for RNA:DNA hybrids in origin-independent replication priming in a eukaryotic system. Proceedings of the National Academy of Sciences of the United States of America. 2015;112(18):5779–84.

32. Lombrana R, Almeida R, Alvarez A, Gomez M. R-loops and initiation of DNA replication in human cells: a missing link? Frontiers in genetics. 2015;6:158.

33. Tran PLT, Pohl TJ, Chen CF, Chan A, Pott S, Zakian VA. PIF1 family DNA helicases suppress R-loop mediated genome instability at tRNA genes. Nature communications. 2017;8:15025.

34. Rippe K, Luke B. TERRA and the state of the telomere. Nature structural & molecular biology. 2015;22(11):853–8.

35. Chedin F. Nascent Connections: R-Loops and Chromatin Patterning. Trends in genetics: TIG. 2016;32(12):828–38.

36. Zhang H, Gan H, Wang Z, Lee JH, Zhou H, Ordog T, et al. RPA Interacts with HIRA and Regulates H3.3 Deposition at Gene Regulatory Elements in Mammalian Cells. Molecular cell. 2017;65(2):272–84.

37. Stirling PC, Hieter P. Canonical DNA Repair Pathways Influence R-Loop-Driven Genome Instability. Journal of molecular biology. 2016.

38. Sollier J, Cimprich KA. Breaking bad: R-loops and genome integrity. Trends in cell biology. 2015.

39. Aguilera A, Garcia-Muse T. R loops: from transcription byproducts to threats to genome stability. Molecular cell. 2012;46(2):115–24.

40. Cerritelli SM, Crouch RJ. Ribonuclease H: the enzymes in eukaryotes. The FEBS journal. 2009;276(6):1494–505.

41. Kobil JH, Campbell AG. Trypanosoma brucei RNase HI requires its divergent spacer subdomain for enzymatic function and its conserved RNA binding motif for nuclear localization. Molecular and biochemical parasitology. 2000;107(1):135–42.

42. Cerritelli SM, Frolova EG, Feng C, Grinberg A, Love PE, Crouch RJ. Failure to produce mitochondrial DNA results in embryonic lethality in Rnaseh1 null mice. Molecular cell. 2003;11(3):807–15.

43. El Hage A, Webb S, Kerr A, Tollervey D. Genome-wide distribution of RNA-DNA hybrids identifies RNase H targets in tRNA genes, retrotransposons and mitochondria. PLoS genetics. 2014;10(10):e1004716.

44. Hutchinson S, Glover L, Horn D. High-resolution analysis of multi-copy variant surface glycoprotein gene expression sites in African trypanosomes. BMC Genomics. 2016;17(1):806.

45. Glover L, Alsford S, Beattie C, Horn D. Deletion of a trypanosome telomere leads to loss of silencing and progressive loss of terminal DNA in the absence of cell cycle arrest. Nucleic Acids Res. 2007;35(3):872–80.

46. Aresta-Branco F, Pimenta S, Figueiredo LM. A transcription-independent epigenetic mechanism is associated with antigenic switching in Trypanosoma brucei. Nucleic Acids Res. 2016;44(7):3131–46.

47. Gan W, Guan Z, Liu J, Gui T, Shen K, Manley JL, et al. R-loop-mediated genomic instability is caused by impairment of replication fork progression. Genes & development. 2011;25(19):2041–56.

48. Helmrich A, Ballarino M, Tora L. Collisions between replication and transcription complexes cause common fragile site instability at the longest human genes. Molecular cell. 2011;44(6):966–77.

49. Tuduri S, Crabbe L, Conti C, Tourriere H, Holtgreve-Grez H, Jauch A, et al. Topoisomerase I suppresses genomic instability by preventing interference between replication and transcription. Nature cell biology. 2009;11(11):1315–24.

50. Gomez-Gonzalez B, Garcia-Rubio M, Bermejo R, Gaillard H, Shirahige K, Marin A, et al. Genome-wide function of THO/TREX in active genes prevents R-loop-dependent replication obstacles. The EMBO journal. 2011;30(15):3106–19.

51. Santos-Pereira JM, Herrero AB, Garcia-Rubio ML, Marin A, Moreno S, Aguilera A. The Npl3 hnRNP prevents R-loop-mediated transcription-replication conflicts and genome instability. Genes & development. 2013;27(22):2445–58.

52. Stork CT, Bocek M, Crossley MP, Sollier J, Sanz LA, Chedin F, et al. Co-transcriptional R-loops are the main cause of estrogen-induced DNA damage. eLife. 2016;5.

53. Huertas P, Aguilera A. Cotranscriptionally formed DNA:RNA hybrids mediate transcription elongation impairment and transcription-associated recombination. Molecular cell. 2003;12(3):711–21.

54. Mischo HE, Gomez-Gonzalez B, Grzechnik P, Rondon AG, Wei W, Steinmetz L, et al. Yeast Sen1 helicase protects the genome from transcription-associated instability. Molecular cell. 2011;41(1):21–32.

55. Aguilera A, Gomez-Gonzalez B. DNA-RNA hybrids: the risks of DNA breakage during transcription. Nature structural & molecular biology. 2017;24(5):439–43.

56. Roy D, Zhang Z, Lu Z, Hsieh CL, Lieber MR. Competition between the RNA transcript and the nontemplate DNA strand during R-loop formation in vitro: a nick can serve as a strong R-loop initiation site. Molecular and cellular biology. 2010;30(1):146–59.

57. Sordet O, Redon CE, Guirouilh-Barbat J, Smith S, Solier S, Douarre C, et al. Ataxia telangiectasia mutated activation by transcription- and topoisomerase I-induced DNA double-strand breaks. EMBO reports. 2009;10(8):887–93.

58. Groh M, Lufino MM, Wade-Martins R, Gromak N. R-loops associated with triplet repeat expansions promote gene silencing in Friedreich ataxia and fragile X syndrome. PLoS genetics. 2014;10(5):e1004318.

59. Britton S, Dernoncourt E, Delteil C, Froment C, Schiltz O, Salles B, et al. DNA damage triggers SAF-A and RNA biogenesis factors exclusion from chromatin coupled to R-loops removal. Nucleic Acids Res. 2014;42(14):9047–62.

60. Li L, Germain DR, Poon HY, Hildebrandt MR, Monckton EA, McDonald D, et al. DEAD Box 1 Facilitates Removal of RNA and Homologous Recombination at DNA Double-Strand Breaks. Molecular and cellular biology. 2016;36(22):2794–810.

61. Ohle C, Tesorero R, Schermann G, Dobrev N, Sinning I, Fischer T. Transient RNA-DNA Hybrids Are Required for Efficient Double-Strand Break Repair. Cell. 2016;167(4):1001–13 e7.

62. Glover L, Horn D. Trypanosomal histone gammaH2A and the DNA damage response. MolBiochemParasitol. 2012;183(1):78–83.

63. Damasceno JD, Obonaga R, Santos EV, Scott A, McCulloch R, Tosi LR. Functional compartmentalization of Rad9 and Hus1 reveals diverse assembly of the 9-1-1 complex components during the DNA damage response in Leishmania. Molecular microbiology. 2016;101(6):1054–68.

64. Siegel TN, Hekstra DR, Cross GA. Analysis of the Trypanosoma brucei cell cycle by quantitative DAPI imaging. Molecular and biochemical parasitology. 2008;160(2):171–4.

65. Sheader K, Vaughan S, Minchin J, Hughes K, Gull K, Rudenko G. Variant surface glycoprotein RNA interference triggers a precytokinesis cell cycle arrest in African trypanosomes. Proceedings of the National Academy of Sciences of the United States of America. 2005;102(24):8716–21.

66. Shah JS, Young JR, Kimmel BE, Iams KP, Williams RO. The 5’ flanking sequence of a Trypanosoma brucei variable surface glycoprotein gene. MolBiochemParasitol. 1987;24(2):163–74.

67. Ohshima K, Kang S, Larson JE, Wells RD. TTA.TAA triplet repeats in plasmids form a non-H bonded structure. JBiolChem. 1996;271(28):16784–91.

68. Pan X, Liao Y, Liu Y, Chang P, Liao L, Yang L, et al. Transcription of AAT^∗^ATT triplet repeats in Escherichia coli is silenced by H-NS and IS1E transposition. PLoS One. 2010;5(12):e14271.

69. Aitcheson N, Talbot S, Shapiro J, Hughes K, Adkin C, Butt T, et al. VSG switching in Trypanosoma brucei: antigenic variation analysed using RNAi in the absence of immune selection. MolMicrobiol. 2005;57(6):1608–22.

70. McCulloch R, Barry JD. A role for RAD51 and homologous recombination in Trypanosoma brucei antigenic variation. Genes & development. 1999;13(21):2875–88.

71. Dobson R, Stockdale C, Lapsley C, Wilkes J, McCulloch R. Interactions among Trypanosoma brucei RAD51 paralogues in DNA repair and antigenic variation. MolMicrobiol. 2011;81(2):434–56.

72. Hartley CL, McCulloch R. Trypanosoma brucei BRCA2 acts in antigenic variation and has undergone a recent expansion in BRC repeat number that is important during homologous recombination. MolMicrobiol. 2008;68(5):1237–51.

73. Sheader K, te VD, Rudenko G. Bloodstream form-specific up-regulation of silent vsg expression sites and procyclin in Trypanosoma brucei after inhibition of DNA synthesis or DNA damage. JBiolChem. 2004;279(14):13363–74.

74. Bhatia V, Barroso SI, Garcia-Rubio ML, Tumini E, Herrera-Moyano E, Aguilera A. BRCA2 prevents R-loop accumulation and associates with TREX-2 mRNA export factor PCID2. Nature. 2014;511(7509):362–5.

75. Bhatia V, Herrera-Moyano E, Aguilera A, Gomez-Gonzalez B. The Role of Replication-Associated Repair Factors on R-Loops. Genes (Basel). 2017;8(7).

76. Castellano-Pozo M, Santos-Pereira JM, Rondon AG, Barroso S, Andujar E, Perez-Alegre M, et al. R loops are linked to histone H3 S10 phosphorylation and chromatin condensation. Molecular cell. 2013;52(4):583–90.

77. Garcia-Pichardo D, Canas JC, Garcia-Rubio ML, Gomez-Gonzalez B, Rondon AG, Aguilera A. Histone Mutants Separate R Loop Formation from Genome Instability Induction. Molecular cell. 2017;66(5):597–609 e5.

78. Wahba L, Gore SK, Koshland D. The homologous recombination machinery modulates the formation of RNA-DNA hybrids and associated chromosome instability. eLife. 2013;2:e00505.

79. Chang EY, Novoa CA, Aristizabal MJ, Coulombe Y, Segovia R, Chaturvedi R, et al. RECQ-like helicases Sgs1 and BLM regulate R-loop-associated genome instability. The Journal of cell biology. 2017.

80. Barry JD, McCulloch R. Antigenic variation in trypanosomes: enhanced phenotypic variation in a eukaryotic parasite. Advances in parasitology. 2001;49:1–70.

81. Kim HS, Cross GA. TOPO3alpha influences antigenic variation by monitoring expression-site-associated VSG switching in Trypanosoma brucei. PLoS Pathog. 2010;6(7):e1000992.

82. Yu K, Chedin F, Hsieh CL, Wilson TE, Lieber MR. R-loops at immunoglobulin class switch regions in the chromosomes of stimulated B cells. Nat Immunol. 2003;4(5):442–51.

83. McCulloch R, Rudenko G, Borst P. Gene conversions mediating antigenic variation in Trypanosoma brucei can occur in variant surface glycoprotein expression sites lacking 70-base-pair repeat sequences. MolCell Biol. 1997;17(2):833–43.

84. Cahoon LA, Seifert HS. Transcription of a cis-acting, noncoding, small RNA is required for pilin antigenic variation in Neisseria gonorrhoeae. PLoS pathogens. 2013;9(1):e1003074.

85. Cahoon LA, Seifert HS. An alternative DNA structure is necessary for pilin antigenic variation in Neisseria gonorrhoeae. Science. 2009;325(5941):764–7.

86. Wachter J, Masters TL, Wachter S, Mason J, Hill SA. pilS loci in Neisseria gonorrhoeae are transcriptionally active. Microbiology. 2015;161(Pt 5):1124–35.

87. Zhang Q, Siegel TN, Martins RM, Wang F, Cao J, Gao Q, et al. Exonuclease-mediated degradation of nascent RNA silences genes linked to severe malaria. Nature. 2014;513(7518):431–5.

88. Amit-Avraham I, Pozner G, Eshar S, Fastman Y, Kolevzon N, Yavin E, et al. Antisense long noncoding RNAs regulate var gene activation in the malaria parasite Plasmodium falciparum. Proceedings of the National Academy of Sciences of the United States of America. 2015;112(9):E982–91.

89. Guizetti J, Barcons-Simon A, Scherf A. Trans-acting GC-rich non-coding RNA at var expression site modulates gene counting in malaria parasite. Nucleic Acids Res. 2016;44(20):9710–8.

90. Alsford S, Horn D. Single-locus targeting constructs for reliable regulated RNAi and transgene expression in Trypanosoma brucei. MolBiochemParasitol. 2008;161(1):76–9.

91. Livak KJ, Schmittgen TD. Analysis of relative gene expression data using real-time quantitative PCR and the 2(-Delta Delta C(T)) Method. Methods. 2001;25(4):402–8.

92. Langmead B, Salzberg SL. Fast gapped-read alignment with Bowtie 2. Nature methods. 2012;9(4):357–9.

93. Li H, Handsaker B, Wysoker A, Fennell T, Ruan J, Homer N, et al. The Sequence Alignment/Map format and SAMtools. Bioinformatics. 2009;25(16):2078–9.

94. Robinson JT, Thorvaldsdottir H, Winckler W, Guttman M, Lander ES, Getz G, et al. Integrative genomics viewer. Nat Biotechnol. 2011;29(1):24–6.

95. Ramirez F, Dundar F, Diehl S, Gruning BA, Manke T. deepTools: a flexible platform for exploring deep-sequencing data. Nucleic Acids Res. 2014;42(Web Server issue):W187–91.

96. Pertea M, Kim D, Pertea GM, Leek JT, Salzberg SL. Transcript-level expression analysis of RNA-seq experiments with HISAT, StringTie and Ballgown. Nature protocols. 2016;11(9):1650–67.

